# Longitudinal Changes in Intracortical Excitability During Ramadan Fasting: A Paired-Pulse Transcranial Magnetic Stimulation Study

**DOI:** 10.64898/2026.05.06.723313

**Authors:** Meeyoung Kim, Ibrahim Mostafa Abuamr, Alham Jehad Ali Al-Sharman, Nabil Saad, Hanan Khalil, Hikmat Hadoush

## Abstract

Ramadan fasting represents a natural model of prolonged daily intermittent fasting associated with metabolic and circadian alterations. This study investigated longitudinal changes in intracortical excitability across pre-, mid-, and post-Ramadan timepoints in healthy adults observing Ramadan fasting.

Thirty fasting participants underwent paired-pulse transcranial magnetic stimulation at three timepoints (pre-, mid-, and post-Ramadan). A non-fasting control group (n = 11) was assessed at pre- and mid-Ramadan. Conditioned motor-evoked potentials were recorded at interstimulus intervals of 2–10 ms and normalized to unconditioned responses. A linear mixed-effects model assessed effects of Timepoint and interstimulus interval (ISI). Secondary outcomes included blood glucose, cognitive performance, sleep duration, and reaction time.

A significant main effect of Timepoint (p < 0.001) indicated longitudinal modulation of intracortical excitability, with increased MEP ratios at mid-Ramadan and partial persistence post-Ramadan. The ISI effect confirmed the inhibition–facilitation gradient (p < 0.001). The Timepoint × ISI interaction was not significant (p = 0.566), indicating a global shift in excitability without ISI-specific modulation. Blood glucose and sleep duration decreased significantly at mid-Ramadan.

Ramadan fasting is associated with a time-dependent increase in intracortical excitability, most appropriately interpreted as a generalized shift rather than selective modulation of inhibitory or facilitatory circuits. These changes occur in the context of concurrent metabolic and sleep alterations and may reflect combined influences of fasting-related metabolic state and reduced sleep duration; however, these factors cannot be disentangled within the present design.

## Introduction

Ramadan fasting constitutes a naturalistic model of prolonged daily intermittent fasting practiced by over one billion Muslims worldwide. During Ramadan, individuals abstain from food and fluid intake from dawn to sunset for approximately 29–30 days, resulting in daily fasting durations of 12–18 hours. This pattern induces predictable metabolic consequences including reduced circulating glucose, altered lipid metabolism, and shifts in insulin and glucagon secretion (Faris et al., 2020; Jahrami et al., 2020; Qasrawi et al., 2017). Concurrently, modifications in sleep architecture, meal timing, and circadian entrainment are well documented (Almeneessier & BaHammam, 2018; BaHammam & Almeneessier, 2020).

The central nervous system is highly sensitive to both metabolic and circadian variation. Glucose represents the primary fuel for neuronal activity, and reductions in its availability alter synaptic transmission and intracortical excitability through effects on GABAergic interneurons, which are particularly metabolically vulnerable (Ali & Lessan, 2024; Lessan et al., 2018; Lessan & Ali, 2019). Circadian phase shifts influence GABAergic and glutamatergic signaling, thereby modulating the principal mechanisms of intracortical inhibition and facilitation (Lang et al., 2011; Ly et al., 2016).

Paired-pulse TMS provides a validated, non-invasive method to assess intracortical inhibitory and facilitatory circuits in vivo. Short-interval intracortical inhibition (SICI), assessed at ISIs of 2–4 ms, is mediated predominantly by GABA*A* receptor activity at cortical interneurons (Kujirai et al., 1993; Rossini et al., 2015). Intracortical facilitation (ICF), measured at ISIs of 8–10 ms, predominantly reflects glutamatergic excitatory interneuronal activity (Chen, 2004; Di Lazzaro et al., 2005; Kujirai et al., 1993). Changes in the MEP ratio—the conditioned MEP amplitude normalized to the unconditioned MEP—thus provide a sensitive index of shifts in the cortical inhibition–excitation balance.

Prior work on metabolic and sleep-related influences on cortical excitability has yielded inconsistent findings, in part due to cross-sectional designs and single time-point assessments (Lang et al., 2011; Ly et al., 2016). Longitudinal TMS studies across Ramadan are currently lacking. The present study aimed to characterize changes in intracortical excitability across pre-, mid-, and post-Ramadan timepoints. We hypothesized that fasting would be associated with reduced SICI and enhanced ICF, reflecting a net shift towards cortical hyperexcitability, most pronounced at mid-Ramadan with partial recovery thereafter.

## Materials and Methods

### Participants

Forty-one healthy young adults were recruited: a fasting group (n = 30; 20 female, 10 male; mean age 19.9 ± 1.4 years; BMI 25.6 ± 6.0 kg/m²) and a non-fasting control group (n = 11; 9 female, 2 male; mean age 19.5 ± 0.9 years; BMI 23.7 ± 1.4 kg/m²). Demographic characteristics are presented in Table 1. Inclusion criteria comprised: age 18–35 years, no neurological or psychiatric disorder, no psychoactive or neuroactive medication, no history of epilepsy, right-hand dominance, and no TMS contraindications. Groups did not differ significantly on any demographic variable at baseline (all *p* > 0.48; Table 1). A participant flowchart is presented in Figure 1.

**Figure 1.**
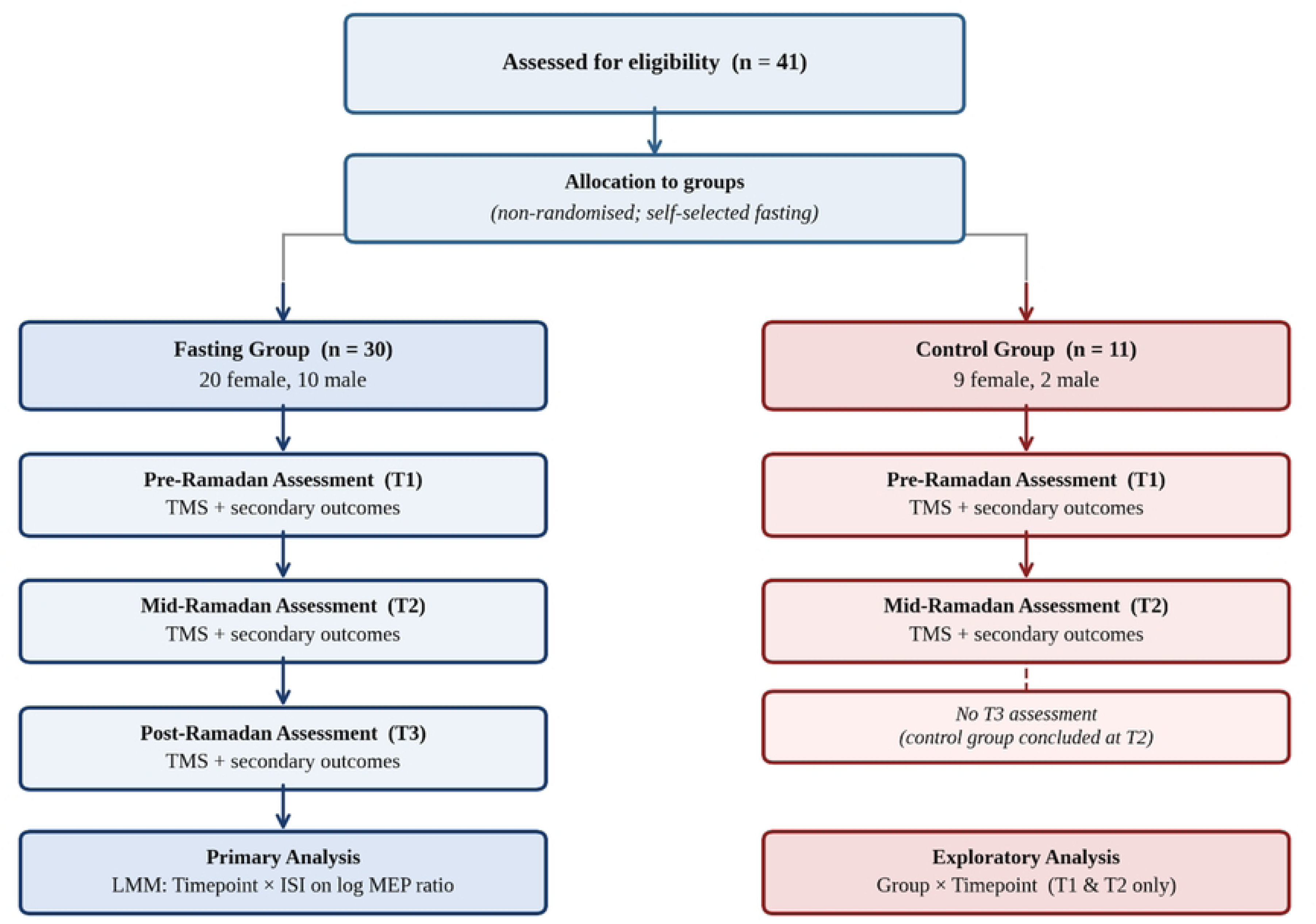
CONSORT-style participant flowchart. Forty-one participants were enrolled following screening. The fasting group (n = 30) was assessed at three timepoints; the control group (n = 11) was assessed at T1 and T2 only. Numbers available for the primary TMS outcome differed from enrolled totals due to missing or unusable MEP data at some ISIs.

**Table 1.**
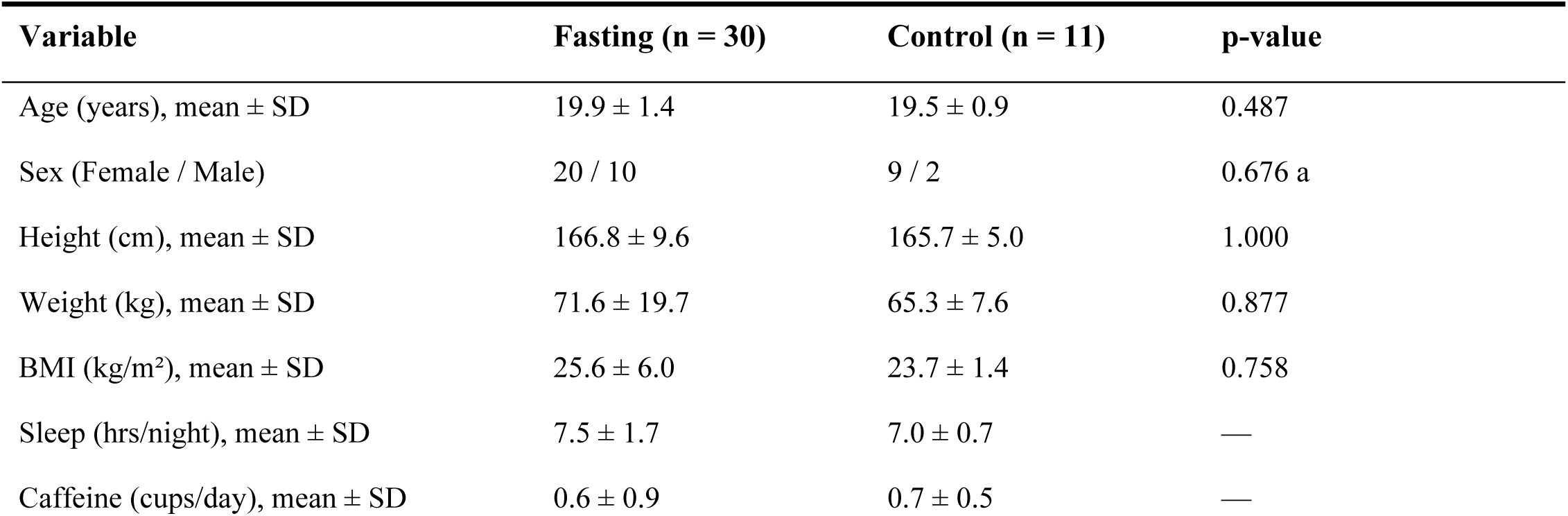
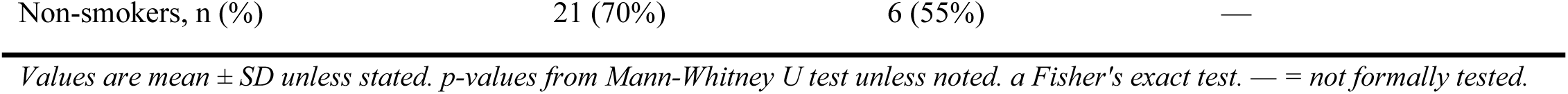
Participant demographic characteristics at baseline.

### Ethics statement

The study was approved by the Research Ethics Unit, University of Sharjah (Reference: REC-25-01-30-01-PG; approved 9 October 2025). Written informed consent was obtained from all participants. Arabic-speaking participants provided consent using an approved Arabic-language consent form.

### Study design and timepoints

A longitudinal repeated-measures design was employed. The fasting group was assessed at T1 (pre-Ramadan, one week before Ramadan), T2 (mid-Ramadan, approximately day 14–16), and T3 (post-Ramadan, during the first week after Ramadan). The control group was assessed at T1 and T2 only. All sessions were conducted between 12:00 and 15:00 to minimize circadian variability in cortical excitability (Lang et al., 2011; Ly et al., 2016). Data collection was conducted between 11 February and 26 March 2026.

### TMS protocol

Cortical excitability was assessed using a DuoMAG MP-Dual stimulator (Deymed Diagnostics, Hronov, Czech Republic) connected to a 70 mm figure-of-eight coil positioned over the hand area of the left primary motor cortex. Surface EMG (TruTrace electromyography) electrodes were placed over the first dorsal interosseous (FDI) muscle of the hand, with a ground electrode positioned over the ipsilateral wrist. The FDI was selected as the target muscle due to its well-characterized cortical representation within M1 and its established use in paired-pulse TMS protocols (Kujirai et al., 1993).

Resting motor threshold (RMT) was defined as the minimum stimulator output required to elicit MEPs of ≥50 μV peak-to-peak amplitude in at least 5 of 10 consecutive trials with the FDI at rest (Rossini et al., 2015). The conditioning stimulus (CS) was set at 80% RMT and the test stimulus (TS) at 120% RMT, consistent with standard paired-pulse protocols (Kujirai et al., 1993; Valls-Solé et al., 1992).

Paired-pulse TMS was administered at ISIs of 2, 4, 6, 8, and 10 ms for the short-interval inhibition block and the rising phase of intracortical facilitation (ICF). A subthreshold conditioning stimulus (80% RMT) preceded suprathreshold TS (120% RMT) at each ISIs. Ten conditioned MEP trials were recorded per ISI, along with ten unconditioned TS trials for normalization. MEP peak-to-peak amplitude (mV) and onset latency (ms) were measured for all trials. All measurements were conducted in the dominant hemisphere only, consistent with established protocols (Kujirai et al., 1993).

### Outcome measures

#### Primary outcome

MEP ratio (conditioned / unconditioned MEP amplitude) for each ISI. Ratios < 1.0 indicate SICI; ratios > 1.0 indicate ICF (Kujirai et al., 1993; Ziemann et al., 1996).

#### Secondary outcomes

Blood glucose (mg/dL; finger-prick glucometry immediately pre-TMS) (Lessan & Ali, 2019), image recall test (Clifford et al., 2024), Stroop color-word task (response time and accuracy) (Periáñez et al., 2021), and simple reaction time (ruler-drop test, cm) (Eckner et al., 2010).

### Statistical analysis

Conditioned MEP amplitudes were divided by the corresponding unconditioned MEP amplitude to yield MEP ratios. Distributions were non-normal (Shapiro-Wilk, all *p* < 0.001); a logarithmic transformation was applied prior to analysis.

A linear mixed-effects model (LMM) assessed fixed effects of Timepoint (3 levels) and ISI (5 levels) and their interaction on log-transformed MEP ratios, with participant as a random intercept. Model significance was evaluated using likelihood ratio tests (LRT) under maximum likelihood estimation (Kujirai et al., 1993; Rossini et al., 2015).

Secondary outcomes were examined using Friedman tests, with Wilcoxon signed-rank post-hoc comparisons (Bonferroni-corrected α = 0.017). Between-group comparisons used the Mann-Whitney *U* test. Effect sizes are reported as rank-biserial *r.* All analyses were conducted in IBM SPSS Statistics (version 29.0; IBM Corp., Armonk, NY, USA). Significance was set at *p* < 0.05 (two-tailed).

### Data availability

All data are publicly available on Zenodo: https://doi.org/10.5281/zenodo.20018903

## Results

### Participant characteristics

Demographic data are in Table 1 and the participant flowchart is in Figure 1. The fasting group comprised 30 participants (20F/10M; age 19.9 ± 1.4 years; BMI 25.6 ± 6.0 kg/m²). The control group comprised 11 participants (9F/2M; age 19.5 ± 0.9 years; BMI 23.7 ± 1.4 kg/m²).

### Unconditioned MEP amplitude and latency

Unconditioned MEP data are presented in Table 2. MEP amplitude showed a non-significant trend in the fasting group (Friedman χ² = 5.08, df = 2, *p* = 0.079). The T1–T3 pairwise comparison showed a medium-effect trend (Wilcoxon *W* = 86, *p* = 0.069, *r* = 0.43) that did not survive the corrected threshold. MEP latency was stable across all three timepoints (Friedman χ² = 2.04, df = 2, *p* = 0.360). No significant between-group differences were observed at T1 or T2 (all Mann-Whitney *p* > 0.05).

**Table 2.**
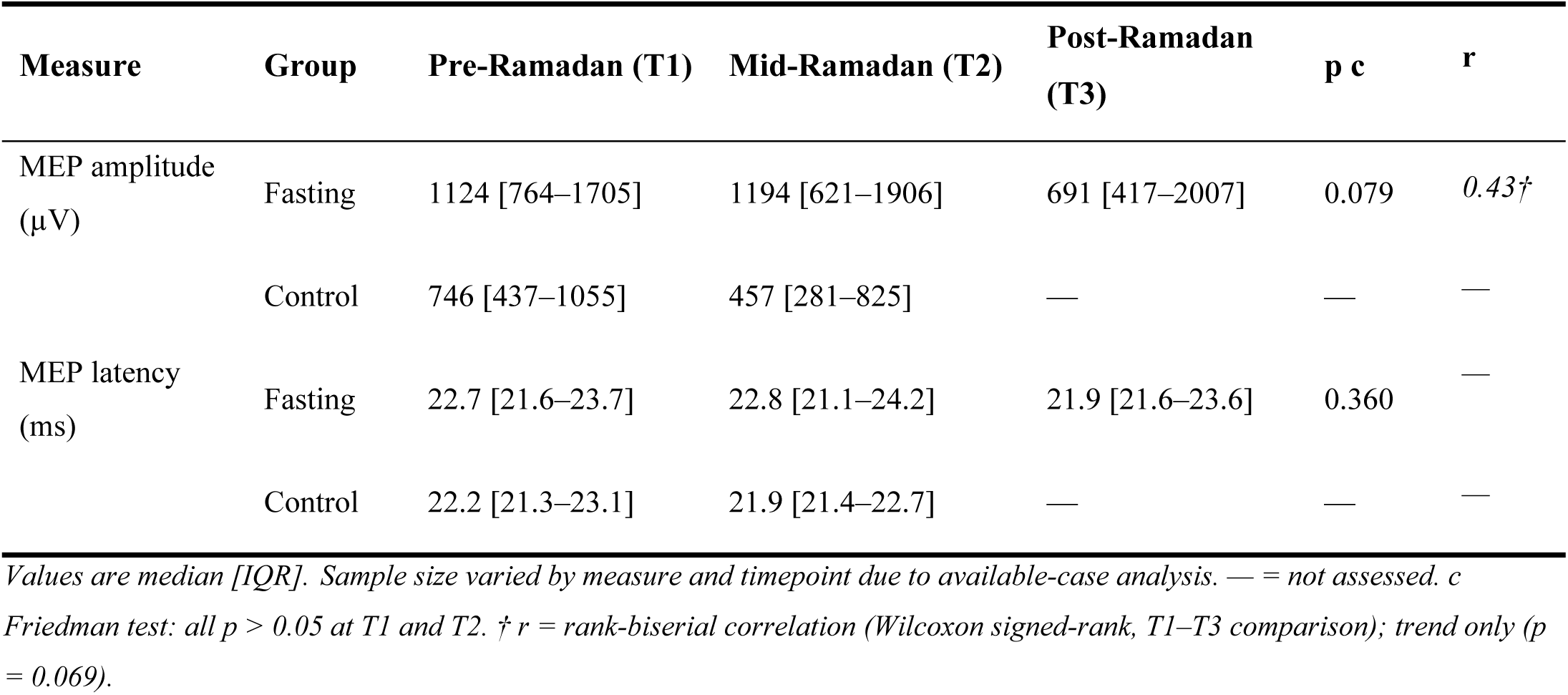
Unconditioned MEP amplitude (µV) and onset latency (ms) by group and timepoint (fasting group; n = 30, control group; n = 11).

### Cortical excitability: paired-pulse TMS

MEP ratio descriptive statistics are in Table 3; LMM results are in Table 4; Figure 2 illustrates the ISI profiles by timepoint. In the fasting group, MEP ratios followed a consistent pattern across all ISIs: lowest at T1, highest at T2, and partially attenuated at T3. The distribution of individual MEP ratios across timepoints is further illustrated in Figure 3.At ISI 2 ms,the mean ratio increased from 0.68 ± 0.51 at T1 to 1.68 ± 2.43 at T2. At ISI 10 ms (ICF), ratios increased from 1.80 ± 1.65 (T1) to 2.98 ± 2.92 (T2), with partial attenuation at T3 (2.59 ± 2.66).

**Figure 2.**
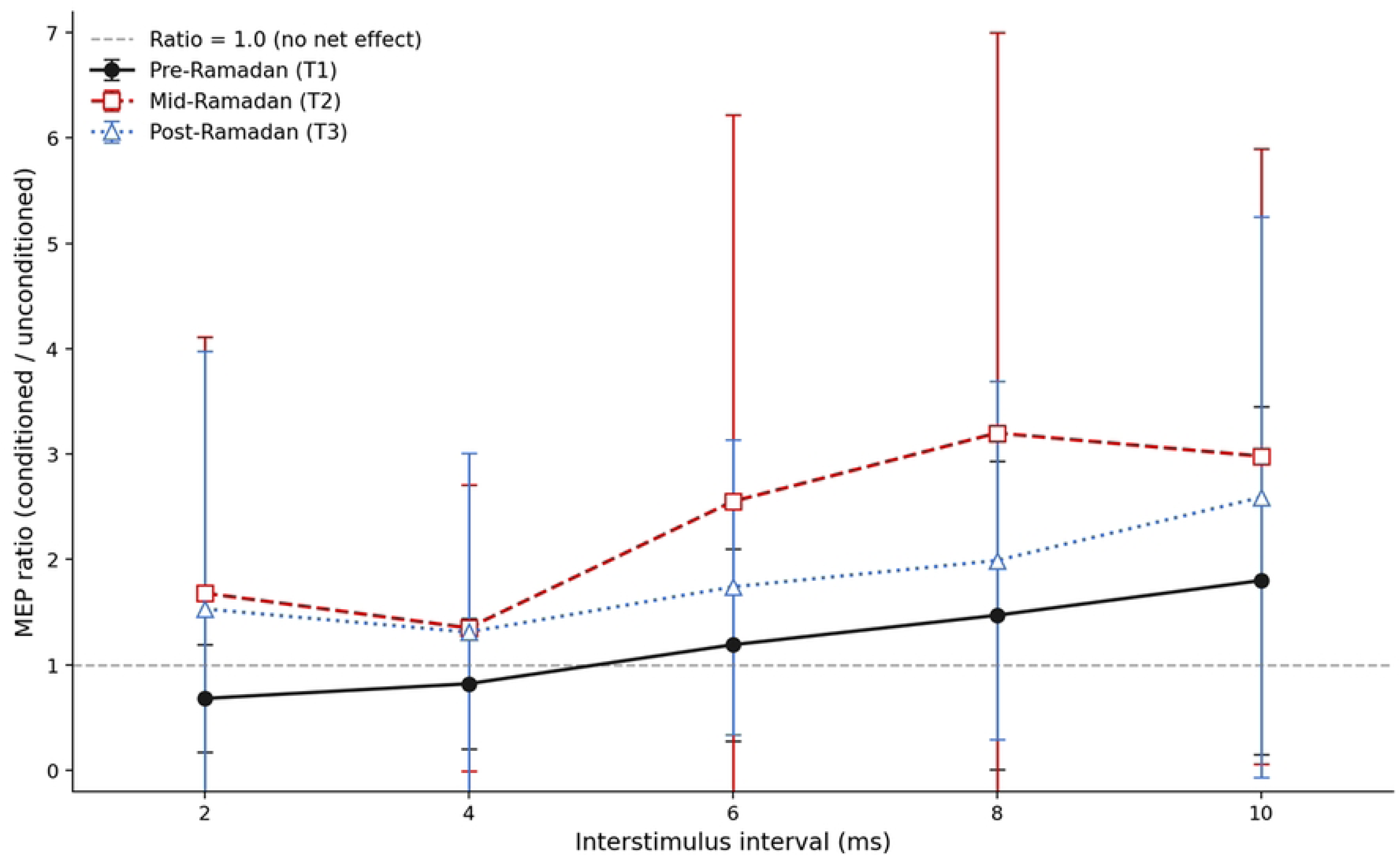
Mean conditioned MEP ratios (± SD) as a function of interstimulus interval (ISI) at each assessment timepoint in the fasting group. MEP ratio = conditioned MEP amplitude / unconditioned MEP amplitude. Values below 1.0 indicate short-interval intracortical inhibition (SICI); values above 1.0 indicate intracortical facilitation (ICF). The horizontal dashed reference line indicates a ratio of 1.0 (no net effect). A significant main effect of Timepoint was observed (χ² = 14.53, df = 2, p < 0.001); the Timepoint × ISI interaction was not significant (p = 0.566).

**Figure 3.**
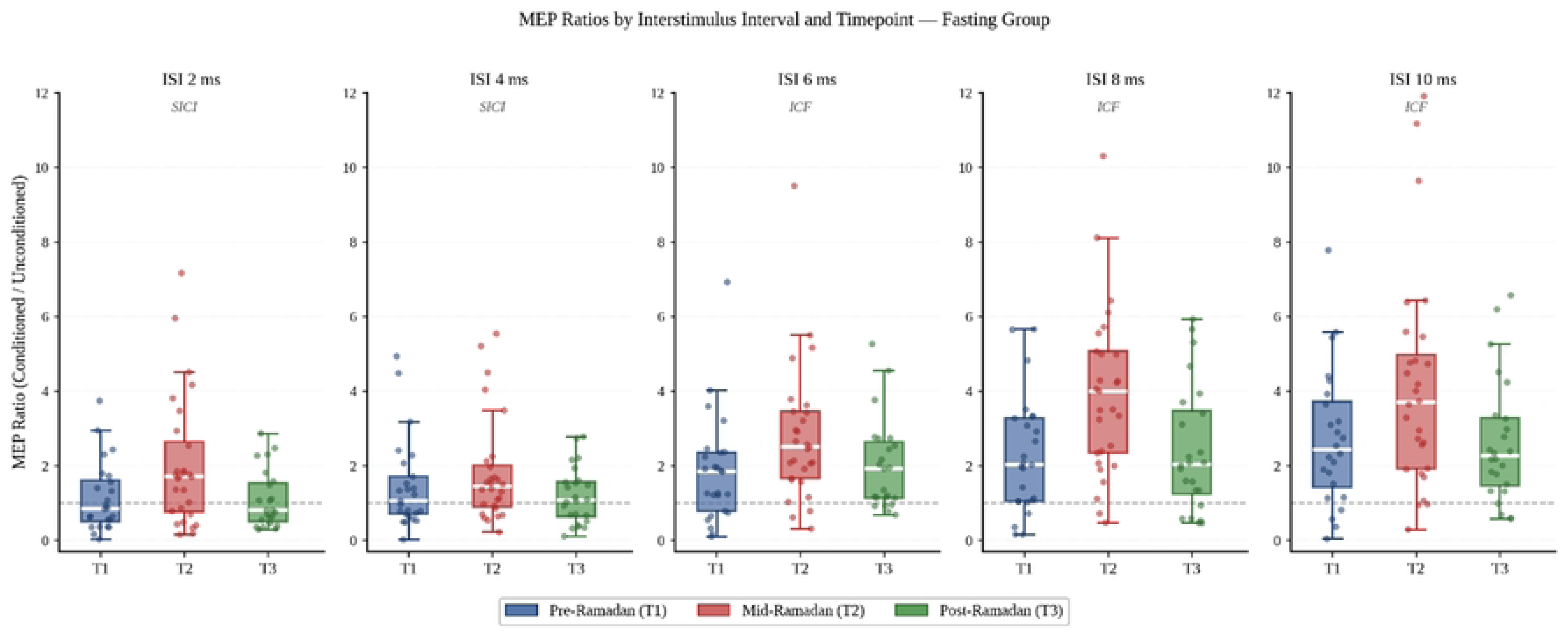
Distribution of conditioned MEP ratios at each interstimulus interval (ISI) across timepoints in the fasting group. Box plots show median (horizontal line) and interquartile range; whiskers extend to 1.5× IQR; values beyond whiskers are not plotted. Individual data points are overlaid with random horizontal jitter for visibility. The horizontal dashed reference line indicates a ratio of 1.0 (no net inhibition or facilitation). Ratios below 1.0 indicate short-interval intracortical inhibition (SICI); ratios above 1.0 indicate intracortical facilitation (ICF). T1 = pre-Ramadan; T2 = mid-Ramadan; T3 = post-Ramadan.

**Table 3.**
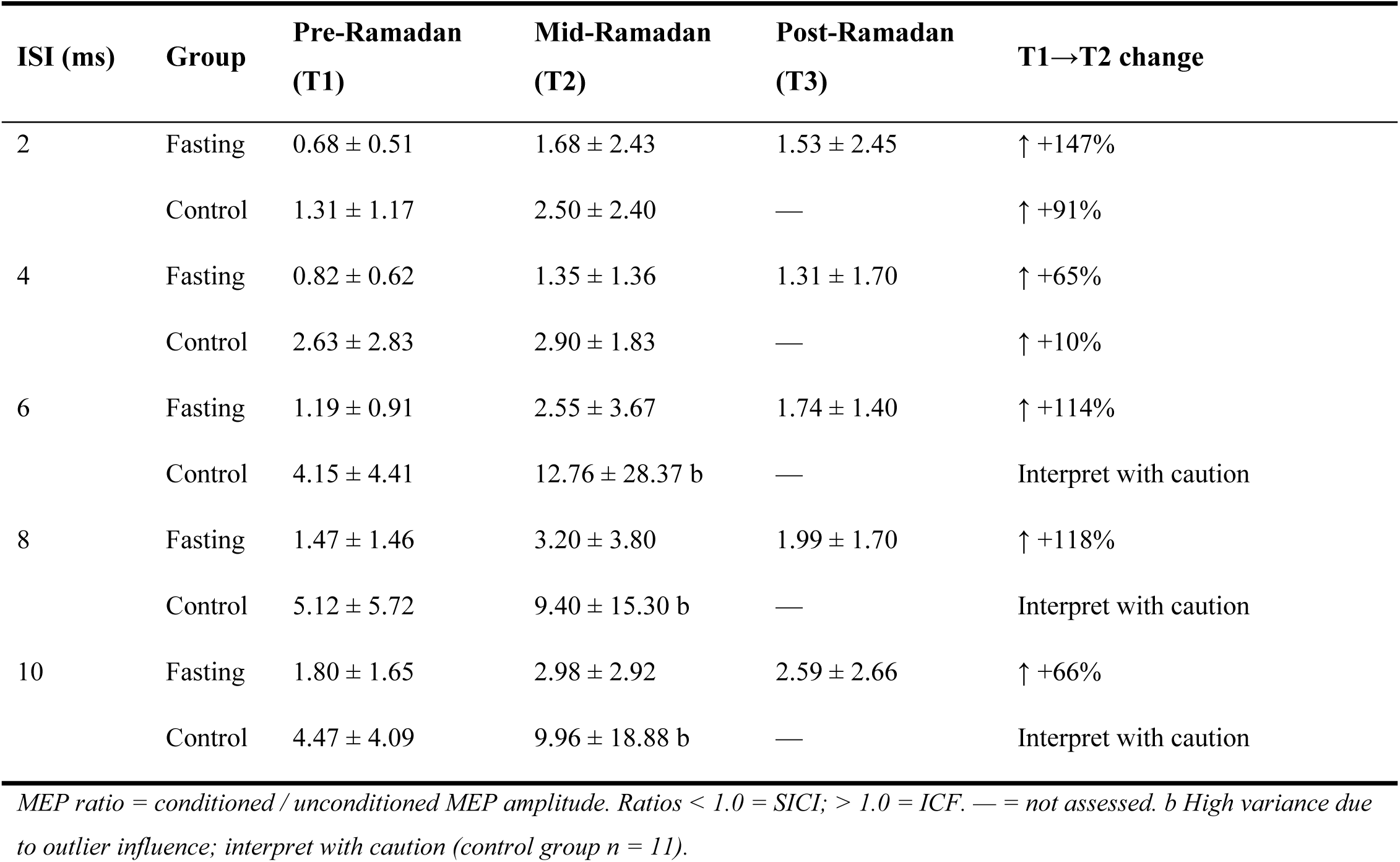
Conditioned MEP ratios (mean ± SD) by ISI, group, and timepoint (fasting group; n = 30, control group; n = 11).

**Table 4.**
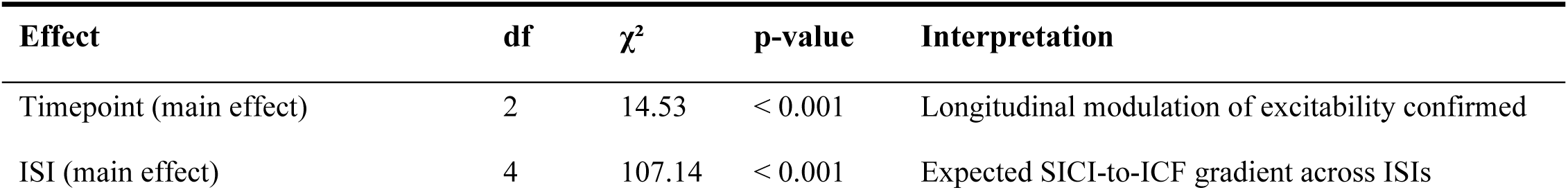
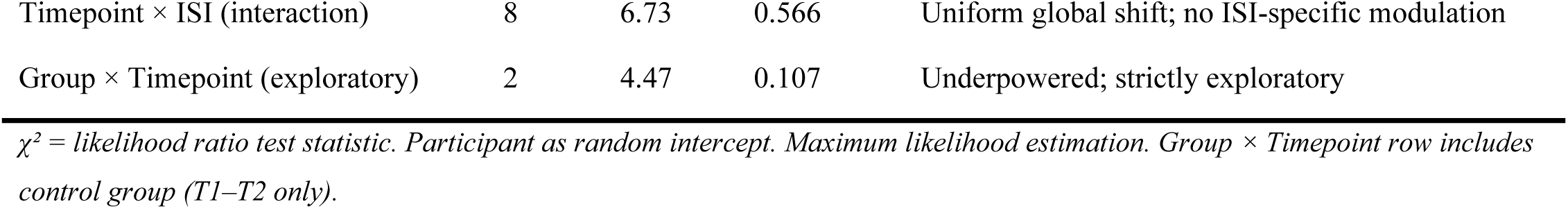
Linear mixed-effects model results (fasting group; n = 30).

The LMM demonstrated a significant main effect of Timepoint (χ² = 14.53, df = 2, *p* < 0.001), and a significant main effect of ISI (χ² = 107.14, df = 4, *p* < 0.001). The Timepoint × ISI interaction was not significant (χ² = 6.73, df = 8, *p* = 0.566). An exploratory analysis including both groups showed a non-significant Group × Timepoint interaction (χ² = 4.47, df = 2, *p* = 0.107).

### Secondary outcomes

Secondary outcome data for both the fasting and control groups are presented in Table 5 and Figure 4. Individual trajectories are shown in Figure 5.

**Figure 4.**
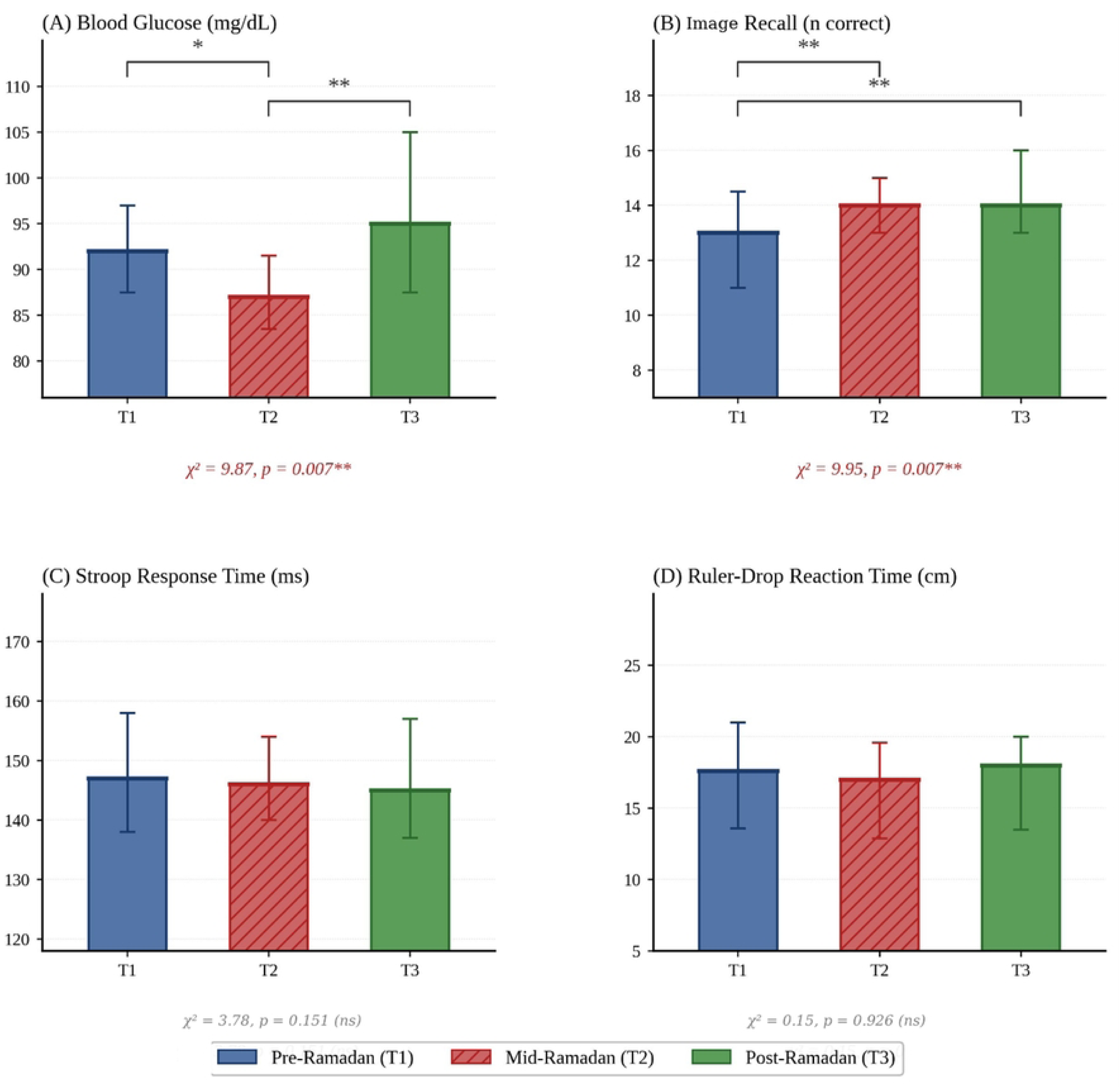
Secondary outcome measures across timepoints in the fasting group (median [IQR]). Bars show medians; error bars denote IQR. Filled bars = Pre-Ramadan (T1); hatched bars = Mid-Ramadan (T2); open bars = Post-Ramadan (T3). Blood glucose decreased significantly at T2 and recovered at T3 (Friedman χ² = 9.87, df = 2, p = 0.007). Image recall improved significantly from T1 to T2 and T3 (Friedman χ² = 9.95, df = 2, p = 0.007). Stroop RT and ruler-drop RT did not change significantly (both p > 0.07). Bonferroni-corrected threshold α = 0.017.

**Figure 5.**
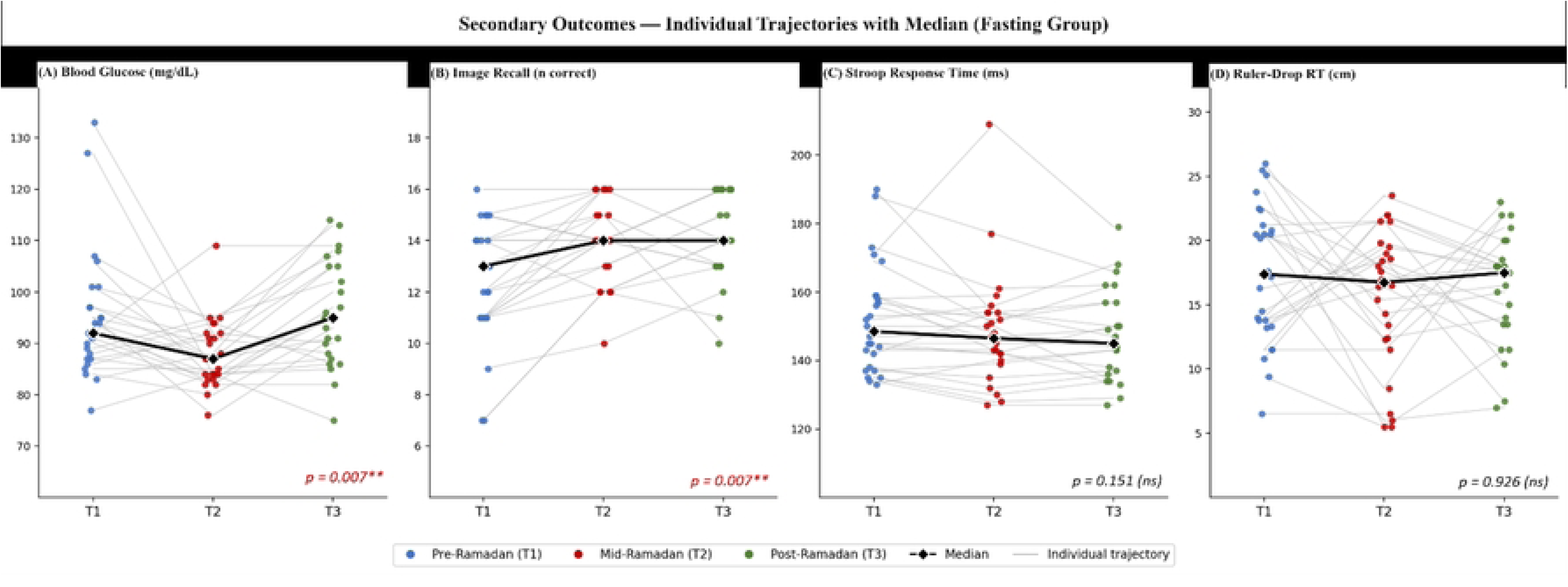
Individual trajectories of secondary outcome measures across timepoints in the fasting group. Grey lines connect each participant’s observations across T1, T2, and T3. Colored dots represent individual data points; vertical-colored bars indicate the interquartile range; the black line with diamond markers represents the group median. (A) Blood glucose (mg/dL); Friedman χ² = 9.87, *p* = 0.007. (B) Image recall (number correct); Friedman χ² = 9.95, *p* = 0.007. (C) Stroop response time (ms); *p* = 0.151 (ns). (D) Ruler-drop reaction time (cm); *p* = 0.926 (ns). Bonferroni-corrected significance threshold α = 0.017. T1 = pre-Ramadan; T2 = mid-Ramadan; T3 = post-Ramadan. ns = not significant.

**Table 5.**
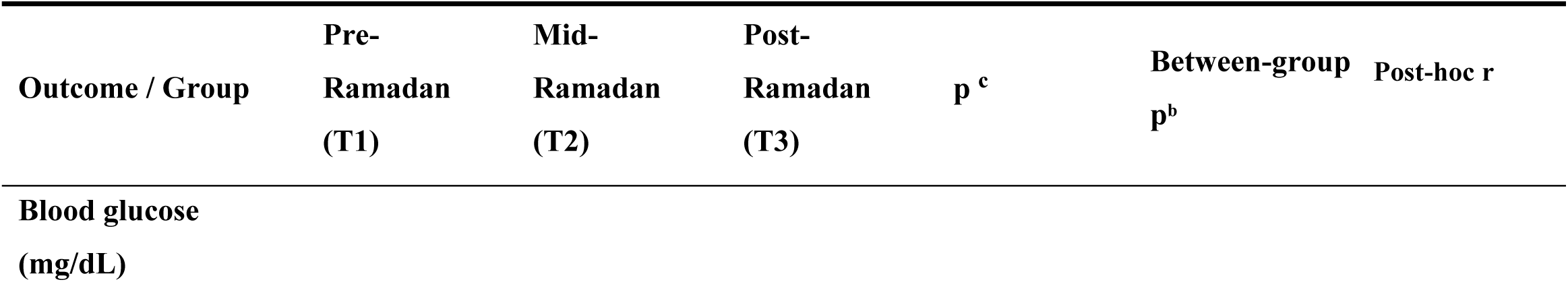
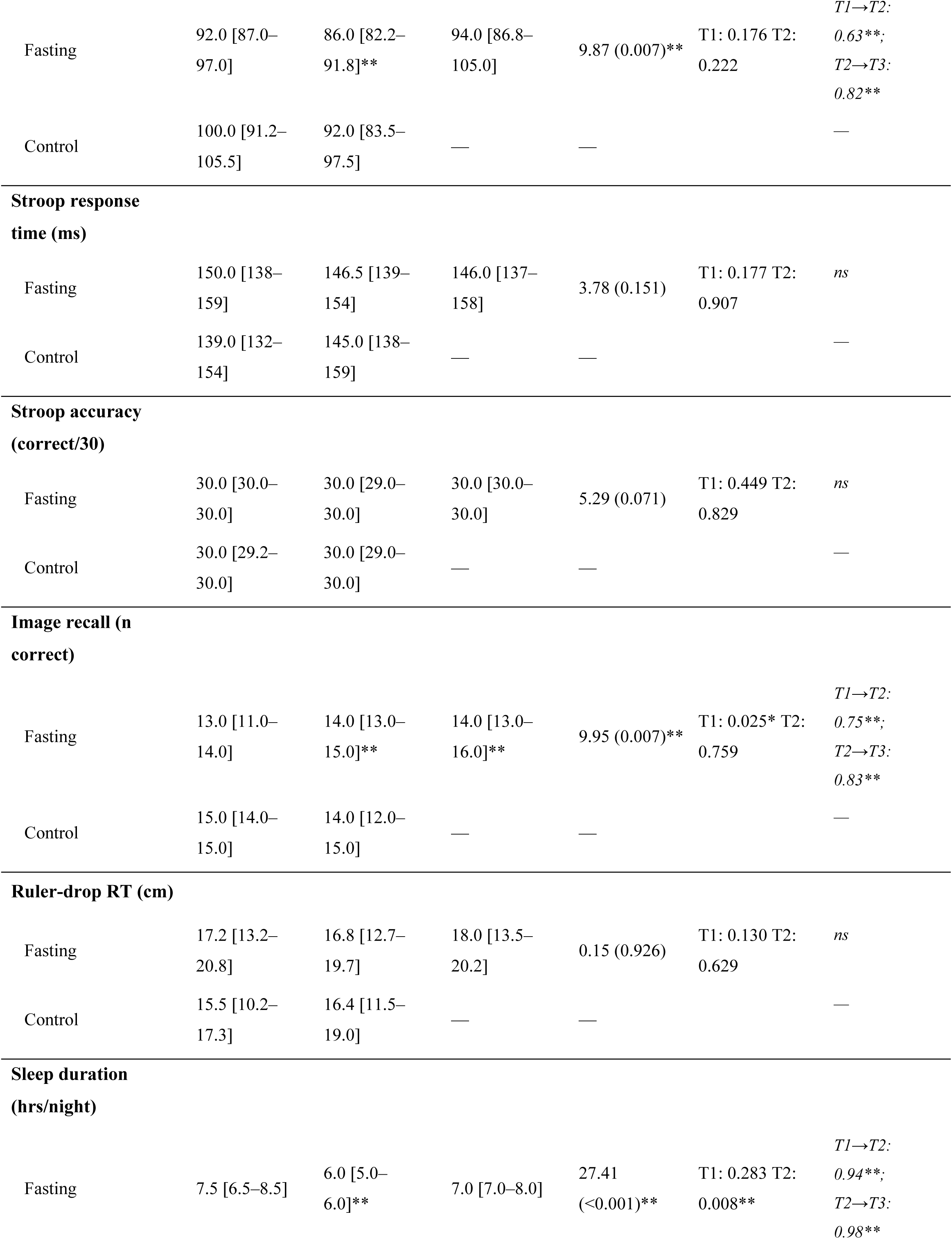
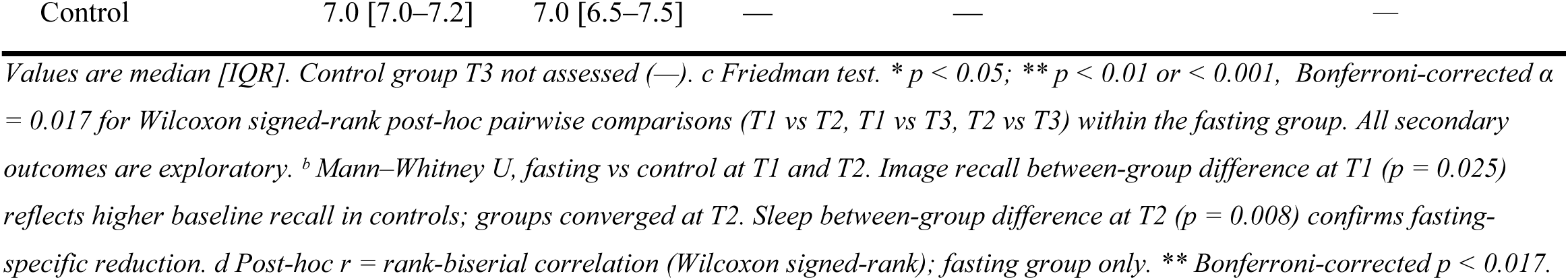
Secondary outcome measures by group and timepoint.

Blood glucose decreased significantly in the fasting group from T1 (median 92.0 mg/dL [IQR 87.0–97.0]) to T2 (86.0 [82.2–91.8]; W = 50, p = 0.014, r = 0.63) and rebounded to T3 (94.0 [86.8–105.0]; W = 25, p = 0.002, r = 0.82). No significant between-group differences were observed at T1 or T2 (p > 0.17).

Image recall increased significantly in the fasting group from T1 (median 13.0 [IQR 11.0–14.0]) to T2 (14.0 [13.0–15.0]; W = 34, p = 0.007, r = 0.75) and T3 (14.0 [13.0–16.0]; W = 24, p = 0.002, r = 0.83). The control group showed higher baseline values at T1 (15.0 [14.0–15.0]) than the fasting group (p = 0.025), with no difference at T2 (p = 0.759).

Stroop response time, Stroop accuracy, and ruler-drop reaction time did not change significantly in the fasting group across timepoints (p > 0.07); these measures did not differ between groups at T1 or T2 (p > 0.13).

Sleep duration decreased significantly in the fasting group from T1 (median 7.5 h [IQR 6.5–8.5]) to T2 (6.0 h [5.0–6.0]; W = 18, p < 0.001, r = 0.94) and recovered at T3 (7.0 h [7.0–8.0]; T2 vs T3: W = 5, p < 0.001, r = 0.98). Sleep duration differed between groups at T2 (p = 0.008).

## Discussion

The present study demonstrates a significant longitudinal modulation of intracortical excitability across the Ramadan period in healthy young adults observing daily intermittent fasting. Conditioned MEP ratios increased from pre-Ramadan (T1) to mid-Ramadan (T2) across all interstimulus intervals (ISIs), with partial attenuation post-Ramadan (T3). These findings indicate a time-dependent shift in intracortical excitability within the fasting group.

A significant main effect of Timepoint was observed in LMM, while the Timepoint × ISI interaction was not significant. This pattern suggests that no ISI-specific modulation was detected. However, the absence of a statistically significant interaction should be interpreted cautiously, as the study may be underpowered to detect differential effects across ISIs. Accordingly, the results are most appropriately interpreted as evidence of a generalized change in MEP ratios over time, rather than selective modulation of inhibitory or facilitatory circuits (Kujirai et al., 1993; Lang et al., 2011; Ziemann et al., 1996).

The observed increase in MEP ratios at shorter ISIs (2–4 ms), alongside increases at longer ISIs (8–10 ms), is consistent with a relative reduction in short-interval intracortical inhibition (SICI) and an increase in intracortical facilitation (ICF) at the group level. These measures are established indices of GABA-A-mediated inhibitory and glutamatergic facilitatory circuits (Kujirai et al., 1993; Rossini et al., 2015; Ziemann et al., 1996). Importantly, unconditioned MEP amplitude and latency remained stable across timepoints, indicating that these changes are unlikely to be attributable to alterations in corticospinal excitability and instead reflect intracortical processes (Fitzgerald et al., 2007; Rothwell, 1997; Ziemann, 2004).

### Intracortical Excitability Changes Across Ramadan

The primary finding of this study — a significant, time-dependent increase in intracortical excitability peaking at mid-Ramadan — extends our understanding of how prolonged intermittent fasting influences cortical inhibition–excitation balance (Lessan & Ali, 2019; Mattson et al., 2017). The pattern of increased MEP ratios across all ISIs at T2, with partial attenuation at T3, suggests that the neurophysiological effects of Ramadan fasting are not static but evolve dynamically across the fasting period and do not fully resolve upon its cessation. This temporal trajectory is consistent with the progressive accumulation of metabolic and circadian perturbations over the course of the month, followed by incomplete physiological recovery in the short post-Ramadan window assessed here (BaHammam & Almeneessier, 2020; Faris et al., 2020).

Ramadan fasting is associated with metabolic and circadian alterations, including changes in glucose availability, sleep timing, and hormonal rhythms (Ajabnoor et al., 2017; Almeneessier & BaHammam, 2018; BaHammam & Almeneessier, 2020; Jahrami et al., 2020). Circadian phase shifts have been shown to modulate cortical excitability and GABAergic inhibition, while metabolic state influences synaptic transmission and neuronal responsiveness (Lang et al., 2011; Ly et al., 2016). These factors provide a plausible physiological context for the observed changes; however, mechanistic interpretations must remain cautious, as the present study did not directly assess neurochemical or circadian markers.

The absence of a significant Timepoint × ISI interaction should be interpreted with caution. While this null result could be taken to suggest a global, ISI-nonspecific shift in excitability, the study was likely underpowered to detect differential modulation across ISIs (Kujirai et al., 1993; Rossini et al., 2015). The observed numerical pattern — with proportionally larger increases at ISI 2 ms and ISI 6 ms — is consistent with a disproportionate reduction in SICI relative to ICF enhancement, which would align with GABAergic impairment as a primary mechanism (Di Lazzaro et al., 2005; Ziemann et al., 1996). However, without adequate statistical power, this pattern cannot be confirmed, and firm conclusions about ISI-specific effects are not warranted (Rossini et al., 2015).

### Metabolic Mechanisms: Glucose and GABAergic Function

Blood glucose levels decreased significantly at mid-Ramadan and rebounded post-Ramadan, consistent with known metabolic adaptations to intermittent fasting (Faris et al., 2019; Lessan & Ali, 2019). Reduced glucose availability has been associated with alterations in neuronal excitability and inhibitory interneuron function, particularly within metabolically demanding GABAergic networks (Alle et al., 2009; Leodori et al., 2019; Toepp et al., 2019). GABAergic interneurons are metabolically demanding and particularly sensitive to reductions in glucose availability. Under conditions of reduced substrate availability, impaired GABAergic tone may lower the threshold for intracortical excitation, manifesting as reduced SICI and relatively enhanced ICF in paired-pulse TMS paradigms (Chen, 2004; Kujirai et al., 1993). This mechanistic account is biologically plausible and is consistent with prior pharmacological work demonstrating that GABA-A receptor modulation produces predictable changes in SICI (Di Lazzaro et al., 2005; Ziemann et al., 1996).

Although this temporal pattern parallels the observed changes in MEP ratios, the study design does not permit causal inference regarding the relationship between metabolic state and intracortical excitability. Blood glucose was measured prior to TMS using finger-prick glucometry, and these values represent a single cross-sectional snapshot rather than a continuous measure of metabolic state during stimulation (Lessan et al., 2018). No direct neurochemical measurements were obtained — plasma or cortical GABA, glutamate, BDNF, cortisol, and insulin were not assayed. Without such measurements, it is impossible to establish whether the observed TMS changes were mechanistically linked to glucose reduction, to downstream GABAergic or glutamatergic changes, or to entirely different metabolic or hormonal pathways (Ali & Lessan, 2024; Lessan & Ali, 2019). Future studies incorporating contemporaneous metabolic and neurochemical measurements at the time of TMS acquisition are required to clarify these relationships. All mechanistic explanations remain speculative working hypotheses, not conclusions supported by the present data.

### Sleep Restriction as a Critical Confound

The concurrent reduction in sleep duration at T2 is a critical confound. In fasting participants, median sleep duration decreased from 7.5 h [IQR 6.5–8.5] at T1 to 6.0 h [5.0–6.0] at T2 (a mean reduction of ∼1.9 h/night; Friedman χ² = 27.41, *p* < 0.001; effect size *r* = 0.94 for T1–T2), recovering fully at T3 (Table 5). Control-group sleep was stable (7.0 h at both T1 and T2) and the between-group difference at T2 was statistically significant (p = 0.008), supporting the interpretation that the sleep reduction was specific to the fasting condition rather than a general seasonal effect.

Sleep restriction independently modulates cortical excitability through GABA-mediated mechanisms (Lang et al., 2011; Ly et al., 2016), and the present design cannot separate fasting-specific effects from those of reduced or shifted sleep. During Ramadan, sleep timing is typically displaced, with delayed sleep onset due to Tarawih prayers and pre-dawn Suhoor meals, resulting in reduced total sleep time and altered sleep architecture (Almeneessier & BaHammam, 2018; BaHammam & Almeneessier, 2020; Faris et al., 2020). The magnitude of sleep reduction observed in this study — approximately 1.9 hours per night — is sufficient, based on prior literature, to plausibly contribute to measurable changes in intracortical excitability (Lang et al., 2011; Ly et al., 2016).

Critically, the T2 TMS session captured participants in a state of concurrent caloric restriction, hydration deprivation, reduced sleep, and circadian phase shift — each of which independently influences cortical excitability through partially overlapping mechanisms (Lessan & Ali, 2019; Ly et al., 2016). This co-occurrence substantially limits causal attribution of the observed MEP ratio changes. Future studies should incorporate polysomnographic or actigraphic sleep monitoring across all timepoints, with statistical control for sleep duration and quality as covariates in the primary analysis (Almeneessier & BaHammam, 2018; Qasrawi et al., 2017).

### Persistence of Elevated Excitability at Post-Ramadan

The persistence of elevated excitability at T3 may reflect residual neurophysiological adaptation following sustained metabolic perturbation (Baik et al., 2020; Ly et al., 2016), though the underlying mechanisms remain unclear. Several non-mutually exclusive explanations may be considered. First, early post-Ramadan may be insufficient for full neurophysiological recovery following 29–30 days of sustained metabolic perturbation; the cortex may undergo homeostatic plasticity changes that outlast the fasting period itself (Baik et al., 2020; Mattson et al., 2017). Second, residual sleep disruption or altered eating patterns in the early post-Ramadan period could sustain excitability elevations beyond the formal end of fasting (Ajabnoor et al., 2017; BaHammam & Almeneessier, 2020). Third, the partial attenuation from T2 to T3 is consistent with a gradual recovery trajectory, suggesting that a longer follow-up period — four to six weeks post-Ramadan — would be needed to establish whether excitability fully normalizes (Fitzgerald et al., 2007; Ly et al., 2016).

It should also be noted that the T3 assessment did not have a matched control group observation, precluding any between-group inference about the post-Ramadan period. Whether the persistent excitability elevation at T3 is specific to fasting individuals, or whether it reflects non-specific seasonal or habituation effects, cannot be determined from the present data (Rossini et al., 2015).

### Cognitive and Functional Correlates

Image recall performance improved significantly in the fasting group across timepoints, with effects persisting post-Ramadan. This observation is consistent with evidence suggesting that intermittent fasting may influence cognitive function and neuroplasticity, potentially through mechanisms involving hippocampal plasticity and neurotrophic signaling pathways (Baik et al., 2020; Seidler & Barrow, 2022). While this pattern is broadly consistent with preclinical evidence suggesting that intermittent fasting may enhance hippocampal neuroplasticity through BDNF-dependent mechanisms (Baik et al., 2020; Mattson et al., 2017; Seidler & Barrow, 2022), several alternative explanations must be considered. Practice effects are a prominent concern: different image sets were used at each timepoint, reducing item-specific practice effects; nonetheless, general task familiarity and response strategy effects across sessions cannot be fully excluded (Zheng et al., 2022) The control group showed numerically stable recall from T1 to T2 (15.0 → 14.0), suggesting no parallel improvement in non-fasting participants, though the control group was too small and was only assessed at two timepoints to draw firm conclusions.

The stability of Stroop response time and ruler-drop reaction time across timepoints suggests that the cognitive effects of Ramadan fasting, if present, may be domain-specific (Eckner et al., 2010; Periáñez et al., 2021). Alternatively, these measures may have been insufficiently sensitive to detect subtle cognitive changes in this healthy young adult sample, where ceiling and floor effects may have attenuated measurable variance (BaHammam et al., 2013; Qasrawi et al., 2017). All cognitive findings are exploratory and should not be interpreted as primary outcomes.

The stability of unconditioned MEP amplitude and latency supports the interpretation that observed ratio changes reflect intracortical rather than corticospinal mechanisms (Fitzgerald et al., 2007; Rothwell, 1997; Ziemann, 2004).

### Limitations and Future Directions

The inclusion of a non-fasting control group provides limited contextual comparison; however, its interpretive value is constrained by small sample size, high variability, and absence of post-Ramadan (T3) assessment. As such, between-group comparisons and interaction analyses are underpowered and should be regarded as exploratory. Additionally, the control group showed higher baseline image recall performance than the fasting group at T1 (p = 0.025), representing a pre-existing between-group difference that limits direct comparison of image recall trajectories across groups. Consequently, the present findings should be interpreted primarily as within-subject longitudinal changes in the fasting cohort, rather than definitive evidence of fasting-specific effects relative to non-fasting individuals.

Several additional limitations should be considered. First, potential confounding variables were not systematically controlled, including sleep duration and timing, hydration status, caffeine intake, and physical activity levels. These factors are known to influence cortical excitability and may vary substantially during Ramadan (Almeneessier & BaHammam, 2018; BaHammam & Almeneessier, 2020; Lang et al., 2011). Second, although testing was conducted within a fixed afternoon window (12:00–15:00), metabolic state at the time of stimulation was not standardized, and time since last meal was not controlled. This represents a limitation, as acute metabolic conditions may directly affect TMS measures (Alle et al., 2009; Toepp et al., 2019).

Third, MEP ratios exhibited substantial within-group variability, particularly at longer ISIs, reflecting the inherent variability of paired-pulse TMS measures (Rossini et al., 2015). Although log transformation was applied to address non-normality, variability may still influence the precision of effect estimates. Larger sample sizes and increased trial counts per condition would improve reliability in future studies.

Finally, the sample comprised predominantly young adults, limiting generalizability to other age groups or clinical populations. The findings therefore should not be extrapolated beyond similar demographic cohorts without further study.

Future studies should include a fully matched control group assessed at all three timepoints, concurrent neurochemical sampling (plasma GABA metabolites, BDNF, cortisol, insulin), continuous actigraphic sleep monitoring, and systematic measurement of hydration and dietary intake (Almeneessier & BaHammam, 2018; Lessan et al., 2018). Pre-specified outlier-handling protocols and adequate statistical power for interaction effects — requiring substantially larger samples than the present study — are required (Rossini et al., 2015). Extending the post-Ramadan follow-up to four to eight weeks would clarify the timescale of cortical recovery (Baik et al., 2020; Ly et al., 2016). Investigation across diverse age groups, fasting practices, and geographical regions would also enhance the generalizability of these findings (Faris et al., 2020; Jahrami et al., 2020).

## Conclusion

Ramadan fasting is associated with a global, time-dependent increase in intracortical excitability, reflected by an overall increase in conditioned MEP ratios across timepoints, peaking at mid-Ramadan and partially persisting after fasting cessation. These changes occur without significant alteration of baseline corticospinal excitability and are accompanied by reductions in blood glucose and alterations in sleep duration.

Given the absence of a significant Timepoint × ISI interaction, the findings are most appropriately interpreted as a generalized shift in intracortical excitability rather than selective modulation of inhibitory or facilitatory circuits. While the observed pattern is consistent with reduced intracortical inhibition and enhanced facilitation, ISI-specific effects were not statistically confirmed.

These results indicate that the metabolic and circadian alterations associated with Ramadan fasting are associated with change in intracortical neurophysiology. However, causal mechanisms remain unresolved due to concurrent changes in sleep and metabolic state and these contributing factors cannot be disentangled within the present design.

## Author Contributions

Conceptualization: MK, IA. Methodology: MK, AJAS, IA. Data curation and formal analysis: MK, IA, HK. Investigation: MK, NS. Writing – original draft: MK, NS. Writing – review and editing: MK, HH, AJAS, HK, IA, NS. Supervision: MK. All authors approved the final manuscript.

## Ethics Statement

The study was conducted in accordance with the Declaration of Helsinki and was approved by the Research Ethics Unit, University of Sharjah (Reference: REC-25-01-30-01-PG; approved 9 October 2025). Written informed consent was obtained from all participants.

## Data Availability

All relevant data are available on Zenodo: https://doi.org/10.5281/zenodo.20018903

## Acknowledgments

The authors would like to acknowledge **Sara Ali Taha** and **Samira K. W. Iqtifan** for their valuable assistance in data collection for this study.

## Use of AI tools

During the preparation of this work, the authors used ChatGPT to assist with language editing and clarity. The authors reviewed and edited all content and take full responsibility for the final manuscript.

## References

1. Ajabnoor, G. M. A., Bahijri, S., Shaik, N. A., Borai, A., Alamoudi, A. A., Al-Aama, J. Y., & Chrousos, G. P. (2017). Ramadan fasting in Saudi Arabia is associated with altered expression of CLOCK, DUSP and IL-1alpha genes, as well as changes in cardiometabolic risk factors. PLOS ONE, 12(4), e0174342. 10.1371/journal.pone.0174342

2. Ali, T., & Lessan, N. (2024). Chrononutrition in the context of Ramadan: Potential implications. Diabetes/Metabolism Research and Reviews, 40(2). 10.1002/dmrr.3728

3. Alle, H., Heidegger, T., Kriváneková, L., & Ziemann, U. (2009). Interactions between short-interval intracortical inhibition and short-latency afferent inhibition in human motor cortex. The Journal of Physiology, 587(21), 5163–5176. 10.1113/jphysiol.2009.179820

4. Almeneessier, A. S., & BaHammam, A. S. (2018). How does diurnal intermittent fasting impact sleep, daytime sleepiness, and markers of the biological clock? Current insights. *Nature and Science of Sleep*, Volume 10, 439–452. 10.2147/NSS.S165637

5. BaHammam, A. S., & Almeneessier, A. S. (2020). Recent Evidence on the Impact of Ramadan Diurnal Intermittent Fasting, Mealtime, and Circadian Rhythm on Cardiometabolic Risk: A Review. Frontiers in Nutrition, 7. 10.3389/fnut.2020.00028

6. BaHammam, A. S., Nashwan, S., Hammad, O., Sharif, M. M., & Pandi-Perumal, S. R. (2013). Objective assessment of drowsiness and reaction time during intermittent Ramadan fasting in young men: a case-crossover study. Behavioral and Brain Functions, 9(1), 32. 10.1186/1744-9081-9-32

7. Baik, S., Rajeev, V., Fann, D. Y., Jo, D., & Arumugam, T. V. (2020). Intermittent fasting increases adult hippocampal neurogenesis. Brain and Behavior, 10(1). 10.1002/brb3.1444

8. Chen, R. (2004). Interactions between inhibitory and excitatory circuits in the human motor cortex. Experimental Brain Research, 154(1), 1–10. 10.1007/s00221-003-1684-1

9. Clifford, J. O., Anand, S., Tarpin-Bernard, F., Bergeron, M. F., Ashford, C. B., Bayley, P. J., & Ashford, J. W. (2024). Episodic memory assessment: effects of sex and age on performance and response time during a continuous recognition task. Frontiers in Human Neuroscience, 18. 10.3389/fnhum.2024.1304221

10. Di Lazzaro, V., Oliviero, A., Saturno, E., Dileone, M., Pilato, F., Nardone, R., Ranieri, F., Musumeci, G., Fiorilla, T., & Tonali, P. (2005). Effects of lorazepam on short latency afferent inhibition and short latency intracortical inhibition in humans. The Journal of Physiology, 564(2), 661–668. 10.1113/jphysiol.2004.061747

11. Eckner, J. T., Kutcher, J. S., & Richardson, J. K. (2010). Pilot Evaluation of a Novel Clinical Test of Reaction Time in National Collegiate Athletic Association Division I Football Players. Journal of Athletic Training, 45(4), 327–332. 10.4085/1062-6050-45.4.327

12. Faris, M. A.-I. E., Jahrami, H. A., Alhayki, F. A., Alkhawaja, N. A., Ali, A. M., Aljeeb, S. H., Abdulghani, I. H., & BaHammam, A. S. (2020). Effect of diurnal fasting on sleep during Ramadan: a systematic review and meta-analysis. Sleep and Breathing, 24(2), 771–782. 10.1007/s11325-019-01986-1

13. Faris, M. A.-I. E., Jahrami, H. A., Obaideen, A. A., & Madkour, M. I. (2019). Impact of diurnal intermittent fasting during Ramadan on inflammatory and oxidative stress markers in healthy people: Systematic review and meta-analysis. Journal of Nutrition & Intermediary Metabolism, 15, 18–26. 10.1016/j.jnim.2018.11.005

14. Fitzgerald, P. B., Fountain, S., Hoy, K., Maller, J., Enticott, P., Laycock, R., Upton, D., & Daskalakis, Z. J. (2007). A comparative study of the effects of repetitive paired transcranial magnetic stimulation on motor cortical excitability. Journal of Neuroscience Methods, 165(2), 265–269. 10.1016/j.jneumeth.2007.06.002

15. Jahrami, H. A., Alsibai, J., Clark, C. C. T., & Faris, M. A.-I. E. (2020). A systematic review, meta-analysis, and meta-regression of the impact of diurnal intermittent fasting during Ramadan on body weight in healthy subjects aged 16 years and above. European Journal of Nutrition, 59(6), 2291–2316. 10.1007/s00394-020-02216-1

16. Kujirai, T., Caramia, M. D., Rothwell, J. C., Day, B. L., Thompson, P. D., Ferbert, A., Wroe, S., Asselman, P., & Marsden, C. D. (1993). Corticocortical inhibition in human motor cortex. The Journal of Physiology, 471(1), 501–519. 10.1113/jphysiol.1993.sp019912

17. Lang, N., Rothkegel, H., Reiber, H., Hasan, A., Sueske, E., Tergau, F., Ehrenreich, H., Wuttke, W., & Paulus, W. (2011). Circadian Modulation of GABA-Mediated Cortical Inhibition. Cerebral Cortex, 21(10), 2299–2306. 10.1093/cercor/bhr003

18. Leodori, G., Thirugnanasambandam, N., Conn, H., Popa, T., Berardelli, A., & Hallett, M. (2019). Intracortical Inhibition and Surround Inhibition in the Motor Cortex: A TMS-EEG Study. Frontiers in Neuroscience, 13. 10.3389/fnins.2019.00612

19. Lessan, N., & Ali, T. (2019). Energy Metabolism and Intermittent Fasting: The Ramadan Perspective. Nutrients, 11(5), 1192. 10.3390/nu11051192

20. Lessan, N., Saadane, I., Alkaf, B., Hambly, C., Buckley, A. J., Finer, N., Speakman, J. R., & Barakat, M. T. (2018). The effects of Ramadan fasting on activity and energy expenditure. The American Journal of Clinical Nutrition, 107(1), 54–61. 10.1093/ajcn/nqx016

21. Ly, J. Q. M., Gaggioni, G., Chellappa, S. L., Papachilleos, S., Brzozowski, A., Borsu, C., Rosanova, M., Sarasso, S., Middleton, B., Luxen, A., Archer, S. N., Phillips, C., Dijk, D.-J., Maquet, P., Massimini, M., & Vandewalle, G. (2016). Circadian regulation of human cortical excitability. Nature Communications, 7(1), 11828. 10.1038/ncomms11828

22. Mattson, M. P., Longo, V. D., & Harvie, M. (2017). Impact of intermittent fasting on health and disease processes. Ageing Research Reviews, 39, 46–58. 10.1016/j.arr.2016.10.005

23. Periáñez, J. A., Lubrini, G., García-Gutiérrez, A., & Ríos-Lago, M. (2021). Construct Validity of the Stroop Color-Word Test: Influence of Speed of Visual Search, Verbal Fluency, Working Memory, Cognitive Flexibility, and Conflict Monitoring. Archives of Clinical Neuropsychology, 36(1), 99–111. 10.1093/arclin/acaa034

24. Qasrawi, S. O., Pandi-Perumal, S. R., & BaHammam, A. S. (2017). The effect of intermittent fasting during Ramadan on sleep, sleepiness, cognitive function, and circadian rhythm. Sleep and Breathing, 21(3), 577–586. 10.1007/s11325-017-1473-x

25. Rossini, P. M., Burke, D., Chen, R., Cohen, L. G., Daskalakis, Z., Di Iorio, R., Di Lazzaro, V., Ferreri, F., Fitzgerald, P. B., George, M. S., Hallett, M., Lefaucheur, J. P., Langguth, B., Matsumoto, H., Miniussi, C., Nitsche, M. A., Pascual-Leone, A., Paulus, W., Rossi, S., … Ziemann, U. (2015). Non-invasive electrical and magnetic stimulation of the brain, spinal cord, roots and peripheral nerves: Basic principles and procedures for routine clinical and research application. An updated report from an I.F.C.N. Committee. Clinical Neurophysiology, 126(6), 1071–1107. 10.1016/j.clinph.2015.02.001

26. Rothwell, J. C. (1997). Techniques and mechanisms of action of transcranial stimulation of the human motor cortex. Journal of Neuroscience Methods, 74(2), 113–122. 10.1016/S0165-0270(97)02242-5

27. Seidler, K., & Barrow, M. (2022). Intermittent fasting and cognitive performance – Targeting BDNF as potential strategy to optimise brain health. Frontiers in Neuroendocrinology, 65, 100971. 10.1016/j.yfrne.2021.100971

28. Toepp, S. L., Turco, C. V., Locke, M. B., Nicolini, C., Ravi, R., & Nelson, A. J. (2019). The Impact of Glucose on Corticospinal and Intracortical Excitability. Brain Sciences, 9(12), 339. 10.3390/brainsci9120339

29. Valls-Solé, J., Pascual-Leone, A., Wassermann, E. M., & Hallett, M. (1992). Human motor evoked responses to paired transcranial magnetic stimuli. Electroencephalography and Clinical Neurophysiology/Evoked Potentials Section, 85(6), 355–364. 10.1016/0168-5597(92)90048-G

30. Zheng, B., Udeh-Momoh, C., Watermeyer, T., de Jager Loots, C. A., Ford, J. K., Robb, C. E., Giannakopoulou, P., Ahmadi-Abhari, S., Baker, S., Novak, G. P., Price, G., & Middleton, L. T. (2022). Practice Effect of Repeated Cognitive Tests Among Older Adults: Associations With Brain Amyloid Pathology and Other Influencing Factors. Frontiers in Aging Neuroscience, 14. 10.3389/fnagi.2022.909614

31. Ziemann, U. (2004). TMS and drugs. Clinical Neurophysiology, 115(8), 1717–1729. 10.1016/j.clinph.2004.03.006

32. Ziemann, U., Rothwell, J. C., & Ridding, M. C. (1996). Interaction between intracortical inhibition and facilitation in human motor cortex. The Journal of Physiology, 496(3), 873–881. 10.1113/jphysiol.1996.sp021734

